## Additional information for "Model guided trait-specific co-expression network estimation as a new perspective for identifying molecular interactions and pathways"

SUPPORTING INFORMATION: S1 APPENDIX

#### Simple numerical examples

**Example 1 (Proposition (1) in the article):** Let us consider an overly simplified interaction model $Y = X_1X_2 + \mathcal{N}(0,1)$. 1000 independent samples of $X_1, X_2 \sim \mathcal{N}(0,1)$ are simulated such that $X_1 \perp\!\!\!\perp X_2$. Truncated correlation matrices $\Sigma^a$ and $\Sigma^{1-a}$ corresponding to the low and high networks are estimated using the truncation point $a = 0.5$. In accordance to the proposition (1) in the article, variables $X_1$ and $X_2$ are negatively correlated in the low group and positively correlated in the high group approximately with the same magnitude:

$$\hat{\Sigma}^a = \begin{bmatrix} 1.00 & -0.423 \\ -0.423 & 1.00 \end{bmatrix} \quad \text{and} \quad \hat{\Sigma}^{1-a} = \begin{bmatrix} 1.00 & 0.425 \\ 0.425 & 1.00 \end{bmatrix}.$$

**Example 2 (Proposition (2) in the article):** Let us now consider an another simple model $Y = \Delta_{1,2}(X_1, X_2) + \mathcal{N}(0,1)$ where $X_1, X_2 \sim \mathcal{N}(0,1)$ and $X_1 \perp\!\!\!\perp X_2$. The type II interaction term $\Delta_{1,2}(X_1, X_2)$ is defined to be $X_1X_2$ if $X_1X_2 \geq 0$ and zero otherwise. As in the previous example, we simulated 1000 independent samples and estimated the high and low networks such that a = 0.5. Now variables $X_1$ and $X_2$ are (positively) correlated only in the high group as can be seen from the estimated correlation matrices below. Respectively, the correlation coefficient between $X_1$ and $X_2$ is nearly zero in the low group:

$$\hat{\Sigma}^a = \begin{bmatrix} 1.00 & -0.04 \\ -0.03 & 1.00 \end{bmatrix} \quad \text{and} \quad \hat{\Sigma}^{1-a} = \begin{bmatrix} 1.00 & 0.86 \\ 0.85 & 1.00 \end{bmatrix}.$$

**Violation of the independence assumption** Let us consider the violation of the assumption (A) in the propositions (1-2). Differential co-expression network type approaches including are poorly capable of finding interaction terms $X_kX_l$ when genes $X_k$ and $X_l$ are strongly correlated before the phenotypic truncation. This is due to fact that the high and low group construction cannot "break" the strong dependency between two genes and therefore the property A1 in the proposition (1) does not hold anymore as shown in the supplementary materials.

**Example:** Let us consider an example were $Y = X_1X_2 + \varepsilon$ without the main effects such that $\varepsilon \sim \mathcal{N}(0, \sigma^2_\varepsilon)$ and $\text{cor}(X_1, X_2) > 0$. This implies that $X_2 = X_1b + \varepsilon_{1|2}$ for some $b \in \mathbb{R}$ and $\varepsilon_{2|1} \sim \mathcal{N}(0, \sigma^2_{\varepsilon_{2|1}})$. With respect to the truncation point $a \in ]0, 0.5]$ we have that

$$\Sigma^a_{1,2} - \Sigma^{1-a}_{1,2} = b\text{var}(X^a_1) + \text{cov}(X^a_1, \varepsilon^a_{1|2}) - b\text{var}(X^{1-a}_1) - \text{cov}(X^{1-a}_1, \varepsilon^{1-a}_{1|2})$$

$$= b(\text{var}(X^a_1) - \text{var}(X^{1-a}_1)) + \text{cov}(X^a_1, \varepsilon^a_{1|2}) - \text{cov}(X^{1-a}_1, \varepsilon^{1-a}_{1|2}).$$

It can be now seen that if the correlation coefficient $b$ is zero, then $\varepsilon^a_{1|2} = X^a_2$ and $\varepsilon^{1-2}_{1|2} = X^{1-a}_2$ so

$$|\Sigma^a_{1,2} - \Sigma^{1-a}_{1,2}| = |\text{cov}(X^a_1, \varepsilon^a_{1|2}) - \text{cov}(X^{1-a}_1, \varepsilon^{1-a}_{1|2})| = |\text{cov}(X^a_1, X^a_2) - \text{cov}(X^{1-a}_1, X^{1-a}_2)| \neq 0.$$

Conversely, when the correlation between variables $X_1$ and $X_2$ becomes close to one, the variance of the error term $\varepsilon_{1|2}$ converges to zero. Then

$$\Sigma^a_{1,2} - \Sigma^{1-a}_{1,2} = \text{var}(X^a_1) - \text{var}(X^{1-a}_1) = 0.$$

Therefore, interaction terms are not identifiable if the correlation between genes is already strong before the phenotypic truncation. The dependencies between genes before truncation should be therefore accounted for by removing these dependencies in the estimation process.

**Example 3 (Violation of the main effect assumption):** Let us consider the following interaction model with two main effects:

$$Y = X_1X_2 + X_1 + X_2 + \varepsilon \quad \text{where} \quad \varepsilon \sim \mathcal{N}(0, 1), \tag{1}$$

where $X_1, X_2 \sim \mathcal{N}(0, 1)$ and $X_1 \perp\!\!\!\perp X_2$. As in the previous example, we simulated 1000 independent samples and estimated the high and low networks such that a = 0.5. In this example, the main effect assumption of the proposition (1) in the article is no longer valid. This implies that the correlations between variables $X_1$ and $X_2$ in the high and low groups are -0.42 and -0.24 (i.e. $\Sigma^a_{1,2} \neq -\Sigma^{1-a}_{1,2}$).

**Example 4 (Residual step):** The previous example continues: If individuals are divided into a high and low groups based on the residual values $\hat{\varepsilon_i}$ estimated from the main effect model, the correlations between variables $X_1$ and $X_2$ in the high and low groups are -0.42 and 0.40 (i.e. $\Sigma^a_{1,2} \approx -\Sigma^{1-a}_{1,2}$). In other words, this additional residual step returns the correlation patterns between variables involved in some important interaction term to behave in accordance to the property A1 even if the main effect assumption of the proposition (1) in the article is not valid.

### Proof for the proposition 1

Here we consider the generalized interaction model [2] (with the scaled response variable) presented and explained in the article:

$$Y_i = \sum_{j=1}^{p} X_{ij}\beta_j + \sum_{k>j} X_{ij}X_{ik}\beta_{jk} + \sum_{k>j} \alpha_{jk}\Delta_{\mathcal{C}_{j,k}}(X_{ij}, X_{ik}) + \varepsilon_i. \tag{2}$$

**Proposition 1** *Let $\Sigma^a_{j,k}$ and $\Sigma^{1-a}_{j,k}$ denote the correlations between variables $X_j$ and $X_k$ in the low and high groups. Variables $X_j$ and $X_k$ are assumed to be independent before the phenotypic truncation (Assumption A) and that the corresponding main effects $\beta_k$ and $\beta_j$ are zero in the model [2] (Assumption B). Then we have that:*

- ***Invariant property:*** *If the interaction term $X_jX_k$ and the response variable $Y$ are independent then*
$$\Sigma^{a}_{j,k} = \Sigma^{1-a}_{j,k} \quad i.e., \quad C_a = |\Sigma^{a}_{j,k} - \Sigma^{1-a}_{j,k}| = 0.$$
- ***Property A1:*** *If the effect $\beta_{jk}$ is non-zero and $\alpha_{jk} = 0$ in the model [2] then $\Sigma^{a}_{j,k} = -\Sigma^{1-a}_{j,k} \neq 0$.*
- ***Property B1:*** *If the effect $\beta_{jk}$ in the model [2] is zero and $\alpha_{jk} > 0$ then $|\Sigma^{a}_{j,k}| > 0$ and $\Sigma^{1-a}_{j,k} = 0$ or vice versa if $\alpha_{jk} < 0$.*

**Proof** Let us first prove the property A1 by rewriting the interaction model [2] as follows:

$$Y = X_jX_k\beta_{jk} + u + \varepsilon.$$

Here $u$ contains all other explanatory terms except $X_j$, $X_k$ and $X_jX_k$. We assume that $u$, $\varepsilon$ (where $u \perp\!\!\!\perp \varepsilon$) and their sum $u + \varepsilon$ are normally distributed (with zero means) and that $X_jX_k \perp\!\!\!\perp u + \varepsilon$. Now by the assumption (A), variables $X_j$ and $X_k$ are independent before the phenotypic truncation. Let us denote the high and low groups with upper indexes $H$ (high) and $L$ (low). For notational convenience, we consider the truncated covariances between variables $X_j$ and $X_k$ instead of correlations:

$$\mathrm{COV}(X_j^H, X_k^H) = E(X_j^H X_k^H) - E(X_j^H)E(X_k^H) \quad \text{and} \quad \mathrm{COV}(X_j^L, X_k^L) = E(X_j^L X_k^L) - E(X_j^L)E(X_k^L). \tag{3}$$

The rest of the proof is based on the theory of truncated regression models – see [1] for introduction. We begin by considering the expected values of the interaction term $X_jX_k$ in truncated samples [1]:

$$E(X_j^H X_k^H) = \int_{\frac{\Phi_Y(1-a)-(u+\varepsilon)}{\beta_{jk}}}^{+\infty} Z_{jk} f_{jk}(Z_{jk}) dZ_{jk} \Bigg/ \left[1 - F_{Z_{jk}}\left(\frac{\Phi_Y(1-a)-(u+\varepsilon)}{\beta_{jk}}\right)\right] \quad \text{and}$$

$$E(X_j^L X_k^L) = \int_{-\infty}^{\frac{\Phi_Y(a)-(u+\varepsilon)}{\beta_{jk}}} Z_{jk} f_{jk}(Z_{jk}) dZ_{jk} \Bigg/ F_{Z_{jk}}\left(\frac{\Phi_Y(a)-(u+\varepsilon)}{\beta_{jk}}\right). \tag{4}$$

Here $f_{jk}(\cdot)$ and $F_{jk}(\cdot)$ are density and cumulative distribution functions for the distribution of the interaction term $Z_{jk} := X_jX_k$ where $f_{jk}(\cdot)$ is assumed to be symmetric and zero-centered, i.e., $f_{jk}(u) = f_{jk}(-u)\, \forall u \in \mathbb{R}$ . Since $E(X_j^H X_k^H)$ and $E(X_j^L X_k^L)$ are functions of the random terms $u$ and $\varepsilon$, we have to evaluate the expected values of $E(X_j^H X_k^H)$ and $E(X_j^L X_k^L)$ with respect to the distribution of the sum $u + \varepsilon$. To that end, let us consider the nominators and denominators of the above ratios [4] separately using shorter notations:

$$\Gamma_H(u{+}\varepsilon, a, \beta_{jk}) = 1 - F_{Z_{jk}}\left(\frac{\Phi_Y(1-a)-(u+\varepsilon)}{\beta_{jk}}\right) \quad \text{and} \quad \Gamma_L(u{+}\varepsilon, a, \beta_{jk}) = F_{Z_{jk}}\left(\frac{\Phi_Y(a)-(u+\varepsilon)}{\beta_{jk}}\right). \tag{5}$$

Symmetricity of the zero-centered density function $f_{jk}(\cdot)$ implies that $\Gamma_H(u{+}\varepsilon, a, \beta_{jk}) = \Gamma_H(-u{-}\varepsilon, a, \beta_{jk})$ since $\Phi_Y(1-a) = -\Phi_Y(a)$. Let us also use shorter notations for the integrals in [4];

$$\Psi_H(u+\varepsilon, a, \beta_{jk}) = \int_{\frac{\Phi_Y(1-a)-(u+\varepsilon)}{\beta_{jk}}}^{+\infty} Z_{jk} f_{jk}(Z_{jk}) dZ_{jk}, \tag{6}$$

$$\Psi_L(u+\varepsilon, a, \beta_{jk}) = \int_{-\infty}^{\frac{\Phi_Y(a)-(u+\varepsilon)}{\beta_{jk}}} Z_{jk} f_{jk}(Z_{jk}) dZ_{jk}. \tag{7}$$

In particular, $\Psi_H(\cdot)$ is a reflection function of $\Psi_L(\cdot)$ i.e., $\Psi_H(u+\varepsilon, a, \beta_{jk}) = -\Psi_L(-u-\varepsilon, a, \beta_{jk})$. It is now straightforward to see that $E(X_j^H X_k^H) = -E(X_j^L X_k^L)$ since

$$\frac{\Psi_H(u+\varepsilon, a, \beta_{jk})}{\Gamma_H(u+\varepsilon, a, \beta_{jk})} = -\frac{\Psi_L(-u-\varepsilon, a, \beta_{jk})}{\Gamma_L(-u-\varepsilon, a, \beta_{jk})} \quad \text{and} \quad E\left(\frac{\Psi_H(u+\varepsilon, a, \beta_{jk})}{\Gamma_H(u+\varepsilon, a, \beta_{jk})}\right) = -E\left(\frac{\Psi_L(-u-\varepsilon, a, \beta_{jk})}{\Gamma_L(-u-\varepsilon, a, \beta_{jk})}\right) \tag{8}$$

due to the symmetric and zero-centered density function of the normally distributed (with a mean of zero) random term $u+e$. Furthermore, the assumptions (A) and (B) together with the distributional assumptions imply that $E(X_j^L) = E(X_k^L) = E(X_j^H) = E(X_k^H) = 0$. Thus

$$\mathrm{COV}(X_j^H, X_k^H) = E\left(\frac{\Psi_H(u+\varepsilon, a, \beta_{jk})}{\Gamma_H(u+\varepsilon, a, \beta_{jk})}\right) = -E\left(\frac{\Psi_L(-u-\varepsilon, a, \beta_{jk})}{\Gamma_L(-u-\varepsilon, a, \beta_{jk})}\right) = -\mathrm{COV}(X_j^L, X_k^L), \tag{9}$$

which are non-zero when $\beta_{jk} \neq 0$. This proves the property A1. The invariant property follows from the assumptions that $X_j X_k \perp\!\!\!\perp (u+\varepsilon)$ and $X_j X_k \perp\!\!\!\perp Y$. The interaction term $X_j X_k$ is therefore independent on the sample selection (based on the response variable) implying that $\Sigma_{j,k}^a = \Sigma_{j,k}^{1-a} = 0$. The property B1 follows immediately from the definition of type II interactions.

### Proof for the proposition 2

**Proposition 2** *Let $\Phi_{j,k}^a$ and $\Phi_{j,k}^{1-a}$ denote the part-correlations between variables $X_j$ and $X_k$ in the low and high groups. Let us assume that the corresponding main effects $\beta_k$ and $\beta_j$ are zero in the model [2]. Then we have that:*

- ***Invariant property:*** *If the interaction term $X_j X_k$ and the response variable $Y$ are independent then $\Phi_{j,k}^a = \Phi_{j,k}^{1-a} = 0$.*
- ***Property C1:*** *If the effect $\beta_{jk}$ is non-zero and $\alpha_{jk} = 0$ in the model [2] then $\Phi_{j,k}^a = -\Phi_{j,k}^{1-a} \neq 0$.*
- ***Property D1:*** *If the effect $\beta_{jk}$ in the model [2] is zero and $\alpha_{jk} > 0$ then $|\Phi_{j,k}^a| > 0$ and $\Phi_{j,k}^{1-a} = 0$ or vice versa if $\alpha_{jk} < 0$.*

**Proof:** See the previous proof for the invariant property. Let us consider the property C1 and assume without loss of generality that $\beta_{jk} = 1$. If $X_j$ and $X_k$ are independent, then the statement follows from the proposition (1). Let us therefore also assume that $X_j$ and $X_k$ are linearly dependent, i.e., $X_j = \rho X_k + \varepsilon_{j|k}$ for some $\rho \in \mathbb{R} \setminus \{0\}$ and $\varepsilon_{j|k} \sim \mathcal{N}(0, \sigma^2_{\varepsilon_{j|k}})$ such that $\varepsilon_{j|k} \perp\!\!\!\perp X_k$. Further, we rewrite the interaction model [2] as follows:

$$Y = X_j X_k \beta_{jk} + u + \varepsilon.$$

Here $u$ contains all other explanatory terms except $X_j$, $X_k$ and $X_j X_k$. We assume that $u$, $\varepsilon$ (where $u \perp\!\!\!\perp \varepsilon$) and their sum $u+\varepsilon$ are normally distributed (with zero means) and that $X_j X_k \perp\!\!\!\perp u + \varepsilon$. These assumptions imply that

$$E(Y) = E(X_j X_k) + E(u+\varepsilon) = E(X_j X_k) = E(\rho X_j^2) + E(X_j \varepsilon_{j|k}) = \rho E(X_j^2).$$

$$\implies E(Y^H) + E(Y^L) = \rho(E(X_j^H)^2 + E(X_j^L)^2) = \rho^H(E(X_j^H)^2) + \rho^L(E(X_j^L)^2),$$

where $\rho^H$ and $\rho^L$ are the regression coefficients in the truncated samples. Due to symmetricity of the zero-centered distribution of $u+\varepsilon$, it can be easily seen that $E(u^H + \varepsilon^H) = -E(u^L + \varepsilon^L)$ using the same reasoning as in the proof of the proposition (1). Therefore,

$$E(Y^H) + E(Y^L) = E(\rho^H (X_j^2)^H) + E(X_j^H \varepsilon_{j|k}^H) + E(\rho^L (X_j^2)^L) + E(X_j^L \varepsilon_{j|k}^L) + E(u^H + \varepsilon^H) + E(u^L + \varepsilon^L)$$

$$\implies E(Y^H)+E(Y^L)-E(\rho^H(X_j^2)^H)-E(\rho^L(X_j^2)^L) = E(X_j^H\varepsilon_{j|k}^H)+E(X_j^L\varepsilon_{j|k}^L)+E(u^H+\varepsilon^H)+E(u^L+\varepsilon^L)$$

$$\implies 0 = E(X_j^H\varepsilon_{j|k}^H) + E(X_j^L\varepsilon_{j|k}^L) + E(u^H+\varepsilon^H) + E(u^L+\varepsilon^L) = E(X_j^H\varepsilon_{j|k}^H) + E(X_j^L\varepsilon_{j|k}^L)$$

$$\implies E(X_j^H\varepsilon_{j|k}^H) = -E(X_j^L\varepsilon_{j|k}^L)$$

which are non-zero when $\beta_{jk} \neq 0$. This proves the property C1. The property D1 follows again immediately from the definition of type II interactions.

### Differential Gaussian graphical model

Here we demonstrate empirically why Gaussian graphical model (GGM) type (conditional dependencies) estimators were not used in our framework. To that end, we generate a simple example in which the simulated response variable is controlled by four different interaction terms such that

$$Y = X_1X_2 + X_1X_3 + X_4X_5 + X_6X_7 + \varepsilon_1 \quad \text{where} \quad \varepsilon_1 \sim \mathcal{N}(0,1),$$

$$\text{and} \quad X_1, X_2, X_4, X_5, X_6, X_7 \overset{\text{i.i.d}}{\sim} \mathcal{N}(0,1).$$

It is further assumed that variable $X_3$ is regulated additively by variables $X_1$ and $X_2$ such that

$$X_3 = X_1 + X_2 + \varepsilon_2 \quad \text{where} \quad \varepsilon_2 \sim \mathcal{N}(0,1). \tag{10}$$

We simulated 1000 independent samples according to this scenario. The dGGM-structure was constructed by the difference of inverse covariance matrices estimated separately for the high and low groups (with the truncation point a = 0.5) with graphical LASSO [2] using the EBIC-criterion for $\lambda-$selection (as expected the smallest possible $\lambda$ were selected). Since the main purpose is to show the difference between the $\Sigma$ and $\Sigma^{-1}$ it is sufficient to use only the dCCN in this section.

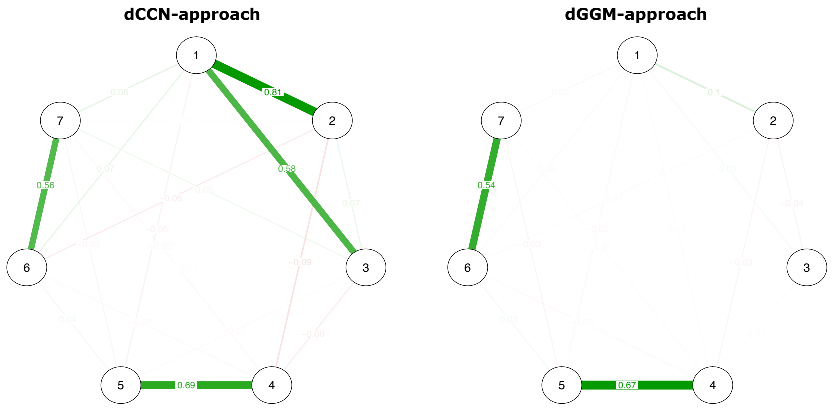


**S1 Fig.** Estimated differential networks with the dCCN and dGGM approaches. Thicknesses of the edges represent the strength of the pairwise dependency between the corresponding genes. The true underlying connections to be found are the edges (1,2), (1,3), (4,5) and (6,7).

It can be seen from S1 Fig that in both cases there are edges between variables associated with the interaction pairs $X_4X_5$ and $X_6X_7$ were each individual variable was simulated independently from other variables. However, the other two true underlying edges (1,2) and (1,3) are visible only in the estimated dCCN structure. This is due to fact that the dependencies between the corresponding variables in the non-truncated dataset interrupt the identifiable patterns characterized in the proposition (1) in the article to occur if conditional dependencies are considered.

#### Fused graphical lasso

Fused graphical LASSO [3] is a method that estimates separate graphical models for multiple observational classes $X^{(k)} \sim \mathcal{N}_p(0, \Sigma_k)$ $(k = 1, 2, \ldots, J)$ and simultaneously controls their structural similarities. It is based on a penalized log-likelihood approach and estimates the precision matrices $\hat{\Theta}^{(k)} = \hat{\Sigma}_k^{-1}$ by solving the following optimization problem

$$\arg\max_{\Theta} \left\{ \sum_{k=1}^{J} n_k \left[ \log(\det(\Theta^{(k)})) - \mathrm{tr}(S^{(k)}\Theta^{(k)}) \right] - P(\Theta) \right\},$$

subject to constraint that $\Theta^{(1)}, \Theta^{(2)}, \ldots, \Theta^{(J)}$ are positive definite matrices. Here $S^{(k)}$ is the sample covariance matrix and the penalty function, denoted by $P(\cdot)$ is designed for encouraging similarity among $\Theta^{(k)}$ and also providing sparsity for each precision matrix. The *fused graphical LASSO* is the solution to the above maximization problem with the penalty function

$$P(\Theta) = \lambda_1 \sum_{k=1}^{2} \sum_{i \neq j} |\Theta_{ij}^{(k)}| + \lambda_2 \sum_{i,j} |\Theta_{ij}^{(2)} - \Theta_{ij}^{(1)}|,$$

where the $\ell_1$-penalty serves to shrink each element $\Theta_{ij}^{(k)}$ towards zero. The second term shrinks the elements of $\Theta^{(1)}, \Theta^{(2)}, \ldots, \Theta^{(J)}$ towards each other. Parameters $\lambda_1$ and $\lambda_2$ in turn are non-negative tuning parameters regulating the amount of shrinkage with respect to the (1) sparsity within each precision matrix ($\lambda_1$) and (2) similarities between them ($\lambda_2$). In the article, we used relatively small values of $\lambda_1$ and $\lambda_2$ due to the hard-thresholding procedure used after the estimation process.
